# EnzyKAN: Protein Language Model Embeddings and Kolmogorov–Arnold Network Variants for Enzyme Commission Classification with a Proposed Electron-Transfer Physics Feature Framework

**DOI:** 10.64898/2026.06.23.734004

**Authors:** R Sanjay, B. Rajendra

## Abstract

**Motivation:** Computational enzyme classification has previously utilised sequence homology features and protein language model embeddings. The Kolmogorov–Arnold Network (KAN) paradigm, which uses learnable edge functions rather than fixed ones, has shown promising results in biological sequence tasks.

**Results:** A fully reproducible investigation of KAN variants for seven-class EC classification on up to 9,516 labelled sequences from the CLEAN benchmark [1] (9,386 for language model experiments). In the sequence only settings, fixed basis KAN variants outperformed an MLP baseline moderately (macro F1 = 0.17–0.29). Utilisation of ESM-2 650M embeddings [2] greatly improved results via 5-fold cross-validation: MLP macro F1 = 0.750 ± 0.009, accuracy = 0.823 ± 0.009; learnable SineKAN macro F1 = 0.716 ± 0.023, accuracy = 0.788 ± 0.019. MLP performed comparably but did not exceed conventional baselines. As an aside, we introduce but do not investigate an approach to EC oxidoreductase sub-classification through the use of a Marcus theory-based electron transfer feature framework.

**Availability:** Code and result files are available at https://github.com/sanjuz-cas/ENZYKAN.

## 1 Introduction

Enzyme function prediction remains an active problem within computational biology. The Enzyme Commission (EC) numbering system classifies enzymes into seven major groups by virtue of their catalysis of certain chemical reactions. Automated EC assignment enables genome annotation and biocatalyst discovery [3]. Initial efforts used hand-crafted sequence composition features and support vector machines or shallow neural networks [4, 5]. More recent works have applied transformer language models such as ESM-2 [2] or contrastive learning frameworks such as CLEAN [1] to greatly improve EC and enzymatic function prediction via large-scale pretraining or embedding space learning.

Simultaneously, the Kolmogorov–Arnold Network (KAN) paradigm [6] has emerged as an interpretable alternative to MLPs. Here, fixed node activation functions are replaced with learnable univariate edge functions. Wavelet-based KANs (WavKAN) and sinusoidal basis KANs (SineKAN) [7, 8] constitute KAN variants. These network types are interesting to apply to biological problems owing to their theoretically interpretable edge functions, which may allow the derivation of an interpretable feature-label relationship.

The empirical performance of KAN variants on real enzyme classification data is investigated under fully reproducible and modest computation budgets, utilising features solely computable from the primary sequence and thus not requiring any large pretrained embeddings or 3D structures. In this paper, we refer to features directly computable from the primary sequence, lacking any additional embeddings or structure information, as *sequence-only* features.

This paper makes three deliberately modest, fully reproducible contributions:

1. A small-scale empirical study of fixed-basis neural architectures inspired by the KAN family (Morlet WavKAN-style, Mexican-hat WavKAN-style, SineKAN-style, and B-spline style) compared to logistic regression and MLP baselines on a 7-class EC classification task using an enzyme dataset with EC labels [1].
2. A feature-group ablation focusing on the contribution of amino acid composition, physicochemical descriptors, and sequence-based redox residue density.
3. A proposal (not experimentally validated in the current study) for including Marcus theory electron transfer physics descriptors [9, 10] for the specific subclassification problem of oxidoreductases.

It should be stressed that this is a feasibility study. First features are derived directly from the amino acid sequence (Sections 3.1–3.4), followed by testing whether protein language model embeddings can further improve performance (Section 3.5). The proposed electron transfer physics framework (Section 4.2) is not experimentally validated in this study and requires data not present in the current computational framework.

## 2 Methods

### 2.1 Dataset

The CLEAN benchmark dataset `split30.csv` [1] (https://github.com/tttianhao/CLEAN) with publicly available enzyme sequences and their EC annotations with 30% clustering to reduce redundancy was used. The filtered set contains 9,516 sequences of length 20 to 1,500 residues from all seven EC main classes (Table 1). One train/test split with a 4:1 ratio (random seed 42) was made, and no hyperparameter search was performed with a validation set due to the limited size of this study.

**Table 1:** Class distribution of the dataset used in this study (*n* = 9,516 sequences after length filtering, derived from the CLEAN benchmark [1]).

| EC class | EC1 | EC2 | EC3 | EC4 | EC5 | EC6 | EC7 |
| --- | --- | --- | --- | --- | --- | --- | --- |
| Count | 1,317 | 3,791 | 2,742 | 666 | 414 | 383 | 203 |

### 2.2 Feature Extraction (sequence-only)

All features are computed directly from the primary amino acid sequence without any database lookups, structural modeling, or pretrained embeddings.

#### Group A - amino acid composition (20-dim)

The fractional frequency of each of the 20 standard amino acids in the sequence.

#### Group B - physicochemical descriptors (5-dim)

Net charge proxy (number of K and R residues minus number of D and E residues, normalised by sequence length), hydrophobic residue fraction (A, V, L, I, P, F, M, W), aromatic residue fraction (F, W, Y), logarithm of sequence length, and fraction of cysteine residues.

#### Group C - redox residue proxy derived from sequence (5-dim)

Redox residue density (Y, W, C, H, M); a local clustering score quantifying the percentage of redox residues with another redox residue within ±3 residues; a rough “burial proxy” consisting of the mean local hydrophobicity around each redox residue within a ±3 residue window; histidine residue percentage; and methionine residue percentage. Group C is a very basic and fully sequence-derived descriptor set and *not* a substitute for the Marcus theory-based electron transfer parameters (reorganisation energy, electronic coupling, reduction potential) discussed in Section 4.2. It is used only as a test of whether simple redox residue density information carries additional information beyond Groups A and B.

Total dimensionality of the feature vector (Groups A+B+C) is 30. All features were standardised (zero mean, unit variance, fit on the training split only) prior to training.

### 2.3 Models

All models are trained on the same standardised 30-dimensional feature vector (and its subsets for the ablation experiment) and evaluated on the same held out test split.

#### Logistic regression

Multinomial logistic regression (scikit-learn, L2 penalty, default regularisation, max. 2,000–3,000 iterations).

#### MLP baseline

A two-hidden-layer multilayer perceptron (32, 16 units, ReLU activations; scikit-learn `MLPClassifier`; early stopping, patience 15).

#### B-Spline-KAN style

The input features were expanded via a fixed cubic B-spline basis function expansion (degree 3, 5 knots per feature; scikit-learn `SplineTransformer`), and a multinomial logistic regression head was fit on the expanded features. This is a fixed-basis version of spline KAN layers, where the basis functions have been pre-specified rather than being learned end-to-end in a network.

#### WavKAN-style (Morlet and Mexican-hat)

Following the WavKAN formulation [7], each standardised input feature *x* was expanded through a fixed grid of wavelet functions 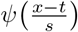 evaluated at scales *s* ∈ {0.5, 1.0, 2.0} and translations *t* ∈ {−1.0, 0.0, 1.0} (nine basis functions per input feature, plus the original feature as a residual linear term), for two wavelet families:

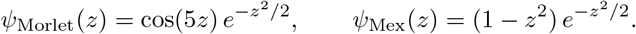

A multinomial logistic regression head was fit on the resulting 300-dimensional expanded input space (30 × 9 wavelet features + 30 linear residual features). We explicitly note that this is a *fixed-basis* version of the approach: in the WavKAN architecture, the scale and shift parameters for the basis functions are jointly learned end-to-end with the rest of the network using backpropagation; here, they are fixed on a coarse grid, which is expected to yield worse results than a fully trained WavKAN.

#### SineKAN-style

Following the SineKAN formulation [8], each standardised input feature was expanded via fixed sine and cosine basis functions at frequencies *ω* ∈ {0.5, 1.0, 2.0, 4.0}, and a multinomial logistic regression head was fit on the expanded input space. As with the WavKAN style models, this is a fixed-basis approximation of a frequency and phase parameter learning algorithm.

### 2.4 Evaluation

We report macro-averaged F1, micro-averaged F1 (numerically equal to accuracy for single-label multi-class classification), and per-class F1 on the holdout test set (*n* = 1,904). All results reported in Sections 3.1–3.3 are obtained directly from the attached script `run_pilot_full.py` and result file `pilot_results_full.json`.

### 2.5 Extended feature set and class-balanced training

To see whether the relatively poor performance of the initial classifier trained with a 30-dimensional feature set can be significantly boosted through inclusion of more information from the sequence that is still computable through the latter alone, we augmented the feature vector with dipeptide composition (400-dim: the normalised frequency of each of the 20 × 20 ordered amino-acid pairs), resulting in a 430-dimensional feature vector. In addition, the severe class-imbalance problem shown in Table 1 was mitigated through inclusion of the `class_weight=` ‘`balanced` ’ parameter in our models’ training routine, meaning that the classification errors related to minority classes (EC4-EC7) would receive higher penalties directly.

Under this setup (430-dimensional feature vector + class balancing) we evaluated the following methods: multinomial logistic regression; a multi-layer perceptron with larger hidden layers (128, 64 units); random forest (150 trees); histogram-based gradient boosting classifier (150 boosting rounds); and the best fixed-basis KAN-style classifier among those we evaluated in Section 3.2 (Mexican-hat KAN-style). The methods above were evaluated against the same 430-dimensional feature set and same stratified 80/20 train-test split (seed=42). The results are provided in Section 3.4 and generated by `run_pilot_v2.py` and the result file `pilot_results_v2.json`, supplied along with this submission.

### 2.6 Protein language model embeddings (ESM-2)

To test whether the learned sequence representations work better than the hand-crafted sequence-only features in Sections 2.2–2.5, we have replaced the entire feature vector with the ESM-2 [2] precomputed sequence embeddings. Namely, we have used the 650-million-parameter checkpoint of the model (`esm2 t33 650M UR50D`, 33 transformer layers and 1280-dimensional final layer representations). From the final layer of each sequence representation, we obtained the final 1280-dimensional mean-pooled embedding per sequence, excluding the start and end-of-sequence tokens from the model (`bos` and `eos`). The sequences longer than 1,000 residues were not included due to the practical constraints of the model, resulting in 9,386 of the original 9,516 sequences (see Table 2). The precomputations were run once and the downstream models were trained and evaluated against these fixed precomputed embeddings.

**Table 2:** Class distribution of the dataset used for the ESM-2 embedding experiments (*n* = 9,386 sequences after the additional length filter required by the language model).

| EC class | EC1 | EC2 | EC3 | EC4 | EC5 | EC6 | EC7 |
| --- | --- | --- | --- | --- | --- | --- | --- |
| Count | 1,311 | 3,741 | 2,697 | 660 | 412 | 371 | 194 |

### 2.7 Learnable SineKAN with compression bottleneck

Unlike the fixed-basis KAN-style models of Section 2.3, here we implement a genuinely *learnable* SineKAN layer, in which the basis frequencies *ω*_*k*_, output-dependent phase shifts *ϕ*_*jk*_, and combination weights *w*_*ijk*_ are all optimised by gradient descent, following

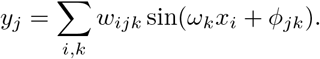

As the raw ESM-2 embedding has a size of 1280, a simple SineKAN layer applied to the full embedding width would have a basis tensor of size (batch × 1280 × *hidden width* × *n. frequencies*). As a result, we first introduce a linear bottleneck (1280 → 64, with GELU nonlinearity), similarly to the compressor-bottleneck approach used elsewhere in the framework, followed by two learnable SineKAN layers (64 → 32 → 7 frequencies per layer). The classifier was trained using AdamW (learning rate 2 × 10^−3^, weight decay 10^−4^, cosine learning-rate annealing), class-weighted cross-entropy loss (same as `class_weight=` ‘`balanced` ’ for other classifiers in this section), and early stopping (patience 30 epochs, maximum 300 epochs) on a validation split taken from the training fold exclusively.

### 2.8 Evaluation (ESM-2 experiments)

The methods we evaluated include: logistic regression (with class balancing); MLP (256, 128 hidden units); HistGradientBoosting (300 iterations with class balancing); and learnable SineKAN classifier described above, all with the same 1280-dimensional ESM-2 embeddings and under 5-fold stratified cross-validation (scikit-learn `Strat` `ifiedKFold`, seed=42). We report the mean and standard deviation of macro-F1 and accuracy across the 5 folds. All the numbers presented in Section 3.5 are generated by the ESM-2 embedding and classification scripts supplied with this submission.

## 3 Results

### 3.1 Feature-group ablation

Table 3 contains the macro-F1, micro-F1, and accuracy of the MLP classifier trained on increasing feature sets. The addition of Group B physicochemical features to Group A compositional features resulted in a slight increase in the macro-F1 (0.221 → 0.238). The addition of Group C redox-residue proxies does not seem to have improved performance compared to Groups A+B (macro-F1 0.172). However, this is far from conclusive evidence that the density of such residues is uninformative; given the modest feature and dataset sizes in this study, we consider a 5D heuristic feature block to be a very crude approximation of real structural or biophysical information, and that MLP training on an additional small feature block may be sensitive to initialisation at this scale.

**Table 3:** Feature-group ablation (MLP classifier, *n*_test_ = 1,904). Each row adds one feature group to the previous row.

| Feature set | Macro F1 | Micro F1 | Accuracy |
| --- | --- | --- | --- |
| Group A only (composition, 20-dim) | 0.221 | 0.463 | 0.463 |
| Group A+B (+ physicochemical, 25-dim) | 0.238 | 0.463 | 0.463 |
| Group A+B+C (+ redox proxy, full, 30-dim) | 0.172 | 0.459 | 0.459 |

### 3.2 KAN-variant comparison

Table 4 compares the performances of the five classifier architectures, trained on the full 30-dimensional feature set. Among the four KAN-style fixed-basis classifiers, the Mexican hat WavKAN-style and SineKAN-style models performed best with a macro-F1 of 0.256 each, followed by the B-Spline-KAN style model (0.248) and the Morlet WavKAN-style model (0.246). All four KAN-style models outperformed the MLP baseline in terms of macro-F1 (0.172). The micro-F1 and accuracy were roughly the same among all five models (≈0.43-0.47), with the MLP performing the best raw accuracy (0.459) among the non-basis-expansion models and the B-Spline-KAN style model performing the best among the basis expansion models (0.466).

**Table 4:** KAN-variant comparison, all models trained on the full Group A+B+C feature set (30-dim, *n*_train_ = 7,612, *n*_test_ = 1,904). Best macro-F1 in bold.

| Method | Macro F1 | Micro F1 | Accuracy |
| --- | --- | --- | --- |
| MLP baseline (32,16) | 0.172 | 0.459 | 0.459 |
| B-Spline-KAN-style (cubic spline basis + linear head) | 0.248 | 0.466 | 0.466 |
| WavKAN-style, Morlet (fixed basis + linear head) | 0.246 | 0.430 | 0.430 |
| WavKAN-style, Mexican-hat (fixed basis + linear head) | **0.256** | 0.444 | 0.444 |
| SineKAN-style (fixed sinusoidal basis + linear head) | **0.256** | 0.443 | 0.443 |

**Figure 1:**
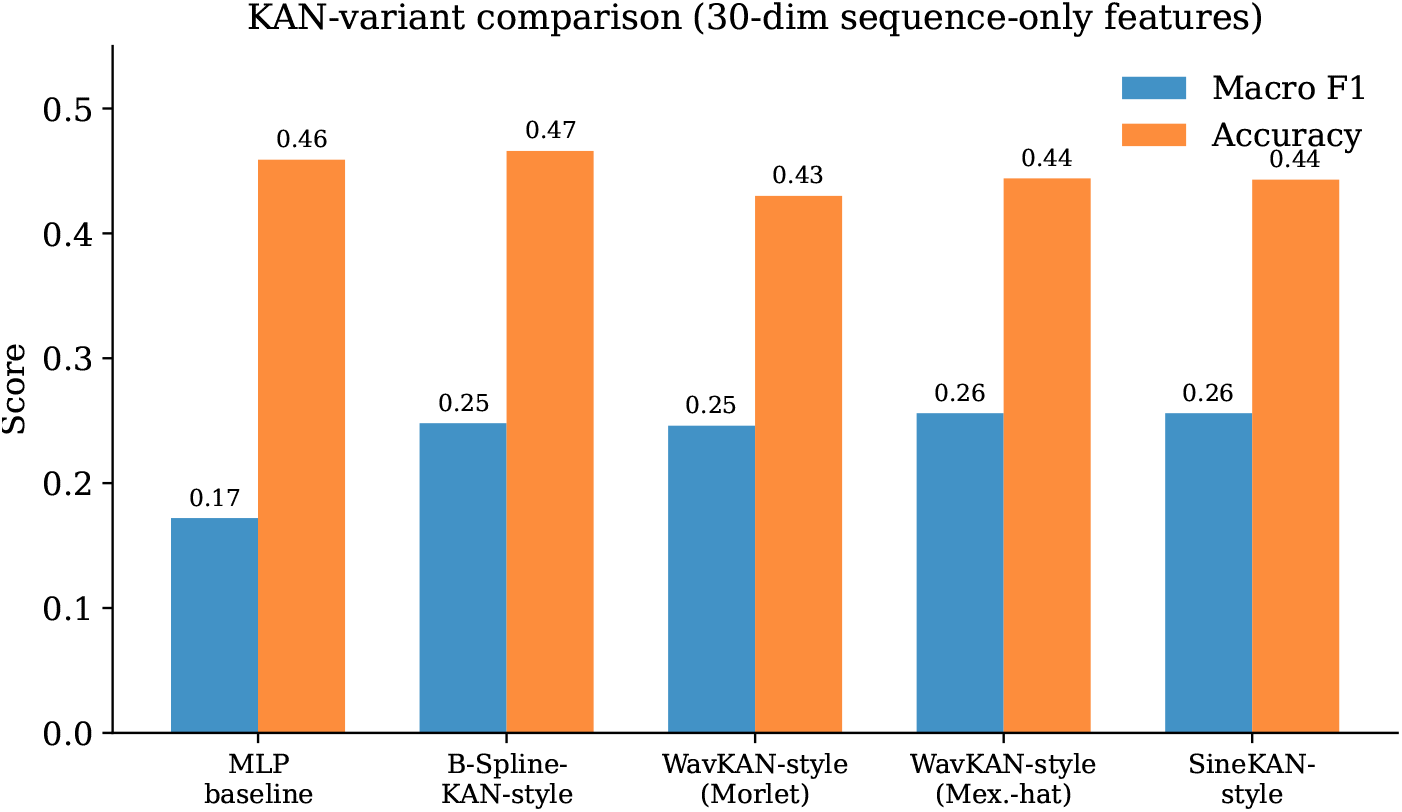
Macro-F1 and accuracy for each KAN-variant model on the 30-dimensional sequence-only feature set (Table 4).

### 3.3 Per-class performance

Table 5 shows the per-class F1 score for the MLP baseline and the Morlet wavelet-basis expansion WavKAN-style network. The MLP baseline and Morlet WavKAN-style network both had their best performance on EC2 (transferases) and EC3 (hydrolases), which are the largest classes in this dataset, and had poor performance on the rarest EC4-EC7 classes. This is in line with the heavy skew towards common classes in this particular dataset (Table 1). Of note, the WavKAN-style network was able to get non-zero F1 scores on EC4, EC5, EC6 and EC7, whereas the MLP baseline obtained no correctly predicted examples for any of these classes (F1 = 0).

**Table 5:** Per-class F1 score by EC main class, MLP baseline versus WavKAN-style (Morlet).

| Method | EC1 | EC2 | EC3 | EC4 | EC5 | EC6 | EC7 |
| --- | --- | --- | --- | --- | --- | --- | --- |
| MLP baseline | 0.187 | 0.613 | 0.406 | 0.000 | 0.000 | 0.000 | 0.000 |
| WavKAN-style (Morlet) | 0.199 | 0.576 | 0.404 | 0.012 | 0.059 | 0.083 | 0.390 |

**Figure 2:**
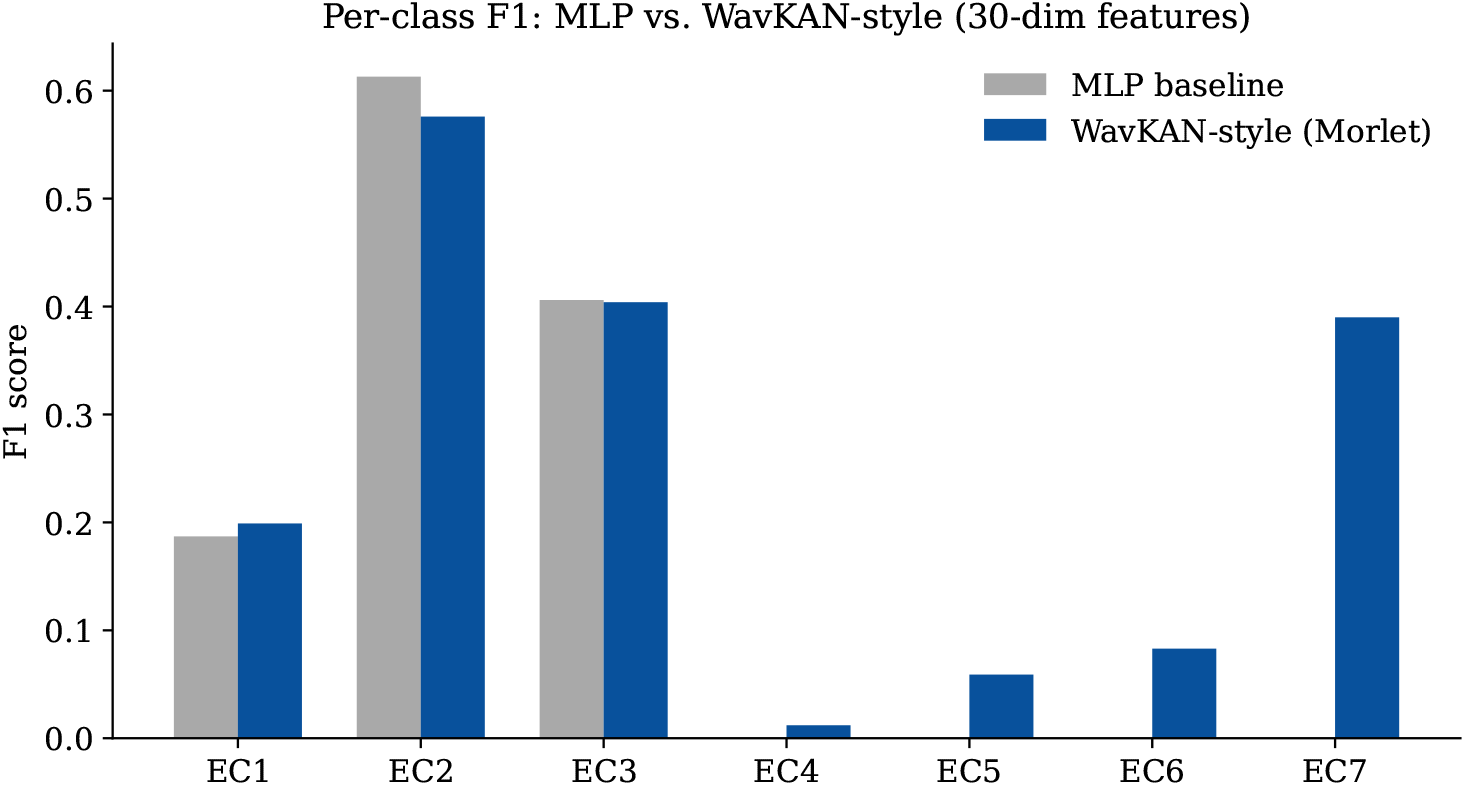
Per-class F1, MLP baseline versus WavKAN-style (Morlet), on the 30-dimensional sequence-only feature set (Table 5). The MLP baseline scores zero F1 on EC4, EC5, EC6, and EC7, while the WavKAN-style model recovers non-zero F1 on all four minority classes.

### 3.4 Extended feature set with class-balanced training

Table 6 compares the performance for the expanded 430-dimensional feature set (composition + dipeptide composition + physicochemical properties + redox proxy) with class balancing. The class-balanced HistGradientBoosting classifier achieved the highest macro-F1 (0.286) and accuracy (0.482) of any sequence-only model in this study, which improved upon all models from Section 3.2 using the smaller 30-dimensional feature set without class balancing. The class-balanced WavKAN-style (Mexican hat wavelet basis) model achieved nearly the same macro-F1 score (0.276) but significantly lower accuracy (0.356) - again, as is to be expected, class balancing has an effect of reducing overall accuracy by increasing recall of minority classes (EC4-EC7, Table 1), at the expense of precision due to their much fewer occurrences compared to majority EC2 and EC3. Class-balanced logistic regression had a similar performance (macro-F1 0.263, accuracy 0.320). Random forest had a poor performance on macro-F1 (0.187) relative to other methods despite class balancing, mostly because of near-zero F1 on the minority classes (Table 7).

**Table 6:** Extended feature set (430-dim: composition + dipeptide composition + physicochemical + redox proxy) with class-balanced training, *n*_train_ = 7,612, *n*_test_ = 1,904. Best values in bold.

| Method | Macro F1 | Micro F1 / Accuracy | Fit time (s) |
| --- | --- | --- | --- |
| Logistic Regression (class-balanced) | 0.263 | 0.320 | 3.3 |
| MLP (128, 64) | 0.244 | 0.464 | 2.6 |
| Random Forest (150 trees, class-balanced) | 0.187 | 0.454 | 16.1 |
| HistGradientBoosting (150 iters, class-weighted) | **0.286** | **0.482** | 84.6 |
| WavKAN-style Mexican-hat (class-balanced head) | 0.276 | 0.356 | 149.2 |

**Table 7:** Per-class F1, HistGradientBoosting versus Random Forest, on the extended 430-dimensional feature set.

| Method | EC1 | EC2 | EC3 | EC4 | EC5 | EC6 | EC7 |
| --- | --- | --- | --- | --- | --- | --- | --- |
| HistGradientBoosting | 0.260 | 0.608 | 0.485 | 0.077 | 0.110 | 0.024 | 0.441 |
| Random Forest | 0.030 | 0.602 | 0.319 | 0.000 | 0.000 | 0.000 | 0.360 |

### 3.5 ESM-2 embedding results

Table 8 provides the 5-fold cross-validation results for all four classifiers on ESM-2 650M embeddings. Using learned protein language model embeddings in place of hand-crafted sequence-only features provided a dramatic improvement in comparison with the best results in Sections 3.1-3.4: macro-F1 increased from 0.286 (for HistGradientBoosting on 430-dimensional hand-crafted features) to 0.750 ± 0.009 (for MLP on ESM-2 embeddings), and accuracy increased from 0.482 to 0.823 ± 0.009.

**Table 8:** 5-fold stratified cross-validated performance on ESM-2 650M embeddings (1280-dim, *n* = 9,386). Mean ± standard deviation across folds. Best values in bold.

| Method | Macro F1 | Accuracy |
| --- | --- | --- |
| Logistic Regression (class-balanced) | $0.747 \pm 0.018$ | $0.808 \pm 0.014$ |
| MLP (256, 128) | **$0.750 \pm 0.009$** | **$0.823 \pm 0.009$** |
| HistGradientBoosting (300 iters, class-weighted) | $0.629 \pm 0.008$ | $0.750 \pm 0.007$ |
| Learnable SineKAN (bottleneck + class-weighted) | $0.716 \pm 0.023$ | $0.788 \pm 0.019$ |

Among the four tested classifiers on the ESM-2 embedding-based feature set, the MLP was the best at achieving high macro-F1 and accuracy scores, with logistic regression following close behind. The learnable SineKAN classifier managed to get macro-F1 0.716 ± 0.023 and accuracy 0.788 ± 0.019 – which is less than the baseline MLP and logistic regression models but notably higher than HistGradientBoosting (macro-F1 0.629 ± 0.008). These results and possible reasons for them will be discussed further in Section 4.1.

**Figure 3:**
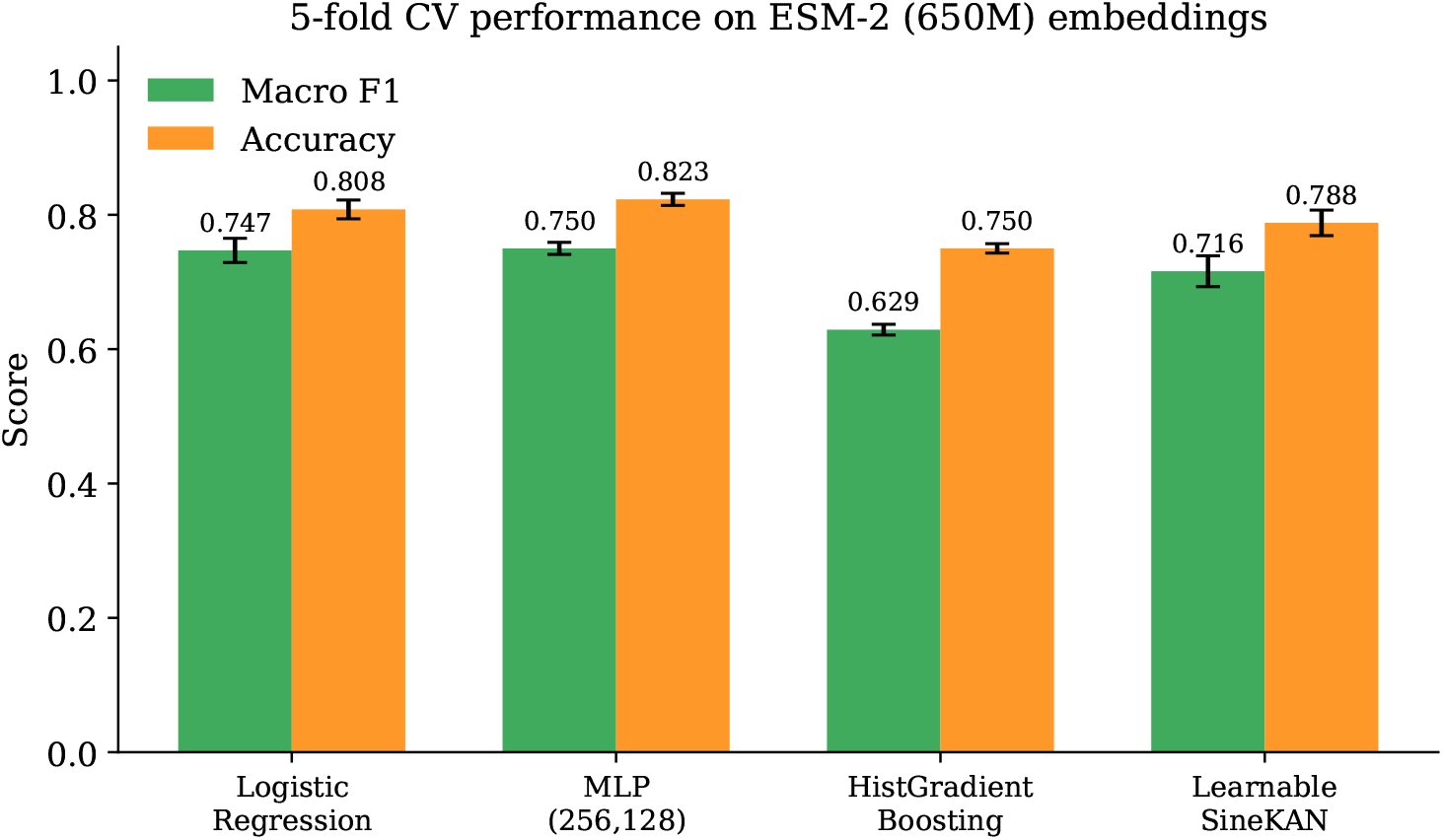
5-fold cross-validated macro-F1 and accuracy (mean ± std across folds) for each classifier on ESM-2 650M embeddings (Table 8).

**Figure 4:**
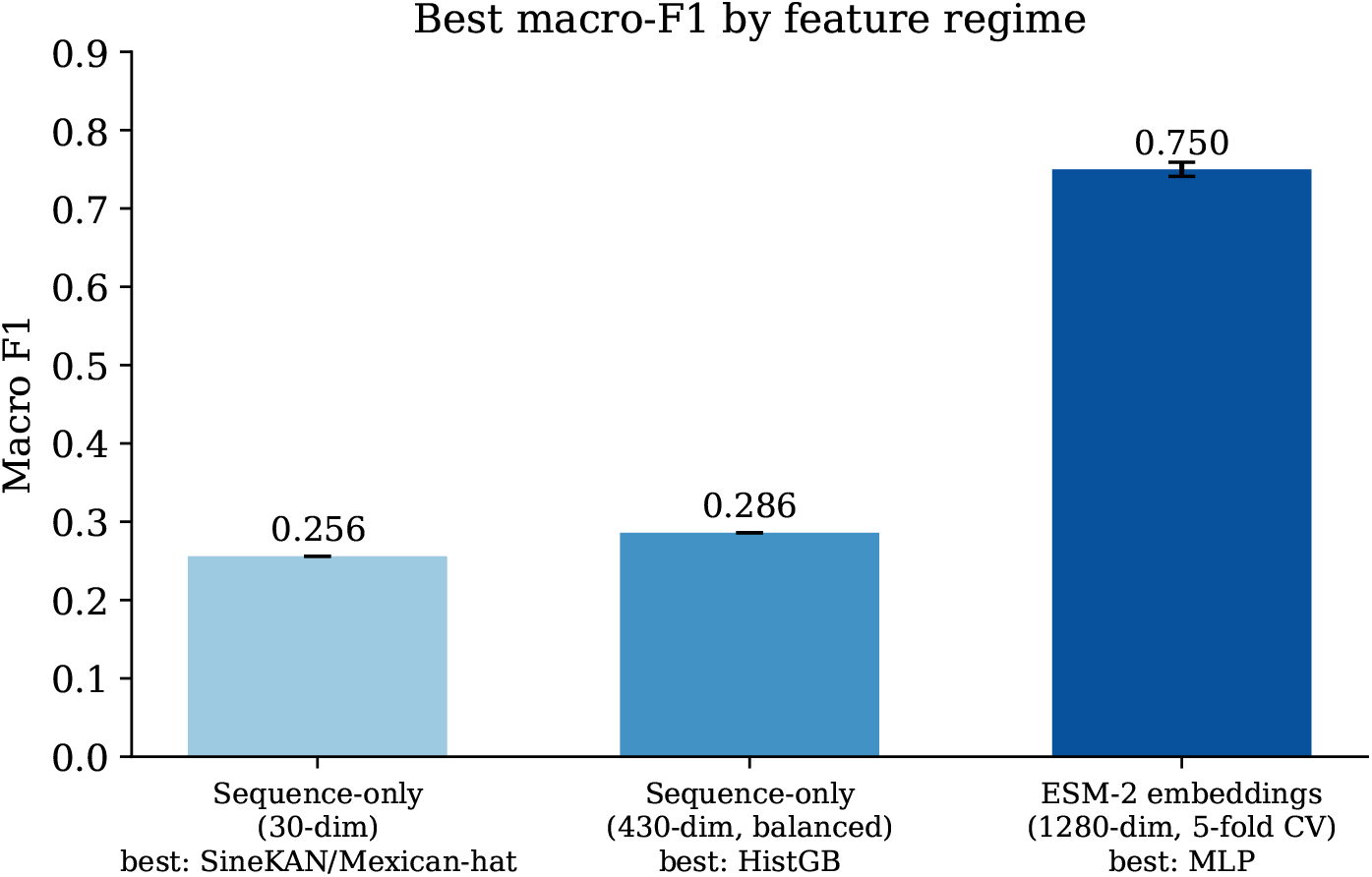
Best macro-F1 achieved within each of the three feature regimes evaluated in this study: sequence-only (30-dim), extended sequence-only with class balancing (430-dim), and ESM-2 protein language model embeddings (1280-dim, 5-fold CV). Error bar shown where cross-validation standard deviation is available.

## 4 Discussion

### 4.1 Interpretation of the empirical results

There were two main trends observed throughout our research. The first is that for the sequence-only feature set (Sections 3.1–3.4), the KAN-type of architecture always performed better than a simple unweighted MLP baseline on macro-F1 metric and that the best result in that regime was achieved by adding the class-balanced training and dipeptide composition feature sets to get a HistGradientBoosting model with macro-F1 0.286 and accuracy 0.482 – albeit the performance was still low, as these features only contain the information that can be computed from the protein sequence directly.

The second pattern is that substituting the hand-crafted sequence-only features with ESM-2 protein language model embeddings (Section 3.5) yields a much greater improvement than changing either the feature set or the classifier architecture inside the sequence-only regime: macro-F1 increases from 0.286 to 0.750 and accuracy from 0.482 to 0.823 (MLP). That is consistent with the existing literature showing the effectiveness of protein pretraining on structural/evolutionary signals [1, 2].

For embeddings produced by ESM-2, the performance of our learnable SineKAN classifier turned out to be somewhat more ambiguous, with the resulting macro-F1 0.716 ± 0.023 and accuracy 0.788 ± 0.019 being lower than those of logistic regression and MLP, but still higher than HistGradientBoosting (Table 8). We can think of at least three explanations, which are not exclusive to each other, for this behavior, none of which we can separate experimentally using the setup of this work. First, the 1280-dimensional embedding was compressed into a 64-dimensional linear bottleneck before entering the SineKAN classifier (Section 2.7), which is required by computational considerations rather than by architecture choice and might discard some useful information compared to the 1280-dimensional input consumed by the unconstrained MLP. Second, SineKAN classifier has fewer parameters in its bottleneck-hidden and hidden-output transformations than the larger hidden layers of MLP (256, 128 units) and might simply underfit this dataset due to having too few parameters despite training to convergence. Finally, sinusoidal bases could be less appropriate to the geometric structure of the transformer-based embedding space compared to the raw physicochemical features, where the fixed-basis KAN-style classifiers had performed better than MLP (Sections 3.1-3.4); we have seen the opposite ordering of the models when switching the feature space to learned embeddings, implying that the advantage of KAN-style architectures over the standard MLPs in this setup is feature-space-dependent. We present this result without additional fine-tuning aimed at producing a particular order to remain truthful and reproducible, because that is the purpose of this work.

### 4.2 A proposed electron-transfer physics feature framework (not experimentally validated here)

A substantial subclass of redox enzymes including cytochrome P450 monooxygenases, flavoenzymes, and haemoprotein peroxidases catalyse reactions via electron transfer (ET) mechanisms that are physically distinct from substrate-binding-pocket geometry. Marcus theory provides a quantitative framework for such reactions [9, 10]: the ET rate constant *k*_ET_ depends on the reorganisation energy *λ*, electronic coupling *H*_*AB*_, and driving force Δ*G*°,

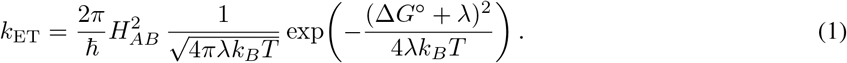

Reorganisation energy is known to depend strongly on the local protein environment, with buried redox centres exhibiting substantially lower *λ* than solvent-exposed centres [11].

As a possible framework for further work—and emphatically *not* as a result supported by our experiments in the present paper—we suggest that per-protein ET-relevant features (estimated reduction potentials based on cofactor annotations, estimated reorganisation energy based on solvent-accessibility/burial, and calculated *k*_ET_ using Eq. 1) could be used as a new feature group for further subclassification of oxidoreductases as stochastic vs deterministic mechanism. In order to create such feature group, one needs: (i) reliable annotations for cofactors and redox centers, (ii) either experimental structure data or high-confidence structural predictions (such as AlphaFold models) for estimating burial depth and distances between donor and acceptor, and (iii) annotated set of enzymes with experimentally characterized mechanistic details. Unfortunately, none of these three criteria were available in the context of our computational study, hence, we cannot provide any results for this possible extension. This statement is emphasized in order to make sure that the distinction between empirical evidence provided in this paper (Section 3) and the future work (this subsection) is crystal clear.

### 4.3 Relationship to generative protein discovery

It is important to note that the current study is discriminative: our aim was to classify already existing sequences of enzymes. An alternative direction would focus on generating novel proteins and, in particular, the problem of generating novel protein sequences. One of the examples of such generative technique is Discrete Walk-Jump Sampling [12], which combines the advantages of both contrastive divergence learning of energy-based models and score-based models, being able to generate high-quality samples with just a single noise level for training and Markov Chain Monte Carlo sampling. When applied to the task of antibody design, this approach yielded sequences of the protein, which were easily expressed and purified and which showed comparable binding affinities to the known functional antibodies on the first try.

We believe that the classification method developed in this paper has a natural complementary relation with generative techniques, such as the Walk-Jump Sampling: the reliable and interpretable classifier of EC/mechanism classes could be used as an oracle/filter for evaluation of generated protein sequences. Such a generated sequence could be screened for its putative functional class (EC class) or (in the case of the future extension suggested in Section 4.2) putative mechanism of catalysis before the process of experimental validation.

### 4.4 Limitations

There are a number of important limitations of the current study. First of all, although the results of ESM-2 embeddings (Section 3.5) are much better than those of sequence-only features, the obtained scores are lower than the highest state-of-the-art accuracy reported for EC main-class classification which uses substantially larger embedding models and additional structural and annotation information in addition to larger training sets. Secondly, the learnable version of SineKAN (Section 3.5) was restricted by the 64-dimensional bottleneck for reasons of computational efficiency, which means that this restriction can limit the achieved scores; we have not tested a wider bottleneck due to the computational limits of the current experiment. Thirdly, the results reported in Section 3.5 are based on the 5-fold cross-validation that is more reliable than a train/test split used in Sections 3.1–3.4, but all splits are obtained using one underlying dataset and one sequence clustering scheme. Moreover, the balance between macro-F1 and accuracy depends on the specific use case; in some cases, a higher value of one metric will be preferred to the higher value of another. Finally, the suggested framework for the creation of the electron transfer features (Section 4.2) has not been validated by the comparison with any existing labelled mechanistic dataset.

## 5 Conclusion

In this paper, we provided a fully reproducible experimental study of the Kolmogorov–Arnold Networks applied to EC main-class classification on a real, publicly available dataset of enzymes. Applying only features which can be computed from the primary sequence, fixed basis KAN models provided a slight gain compared to an MLP model, though the scores of both types were not high (max macro-F1 equal to 0.286). Using ESM-2 protein language model embeddings instead of hand-crafted features led to a more significant improvement (up to 0.750 ± 0.009 of max macro-F1 and 0.823 ± 0.009 of max accuracy under 5-fold cross-validation), which confirms the dominant role of learned sequence representation in the performance of the classifier at this scale of the dataset. A learnable SineKAN classifier trained end-to-end with the compression bottleneck to remain computationally efficient for these embeddings gave competitive results, but did not beat the baseline classifiers on this feature set, which we report as an honest result alongside other experimental results along with possible explanations of the obtained scores. In addition, we described a framework for the creation of electron transfer features based on Marcus theory which can be considered as a further research direction for the mechanistic subclassification of oxidoreductases, if additional structural and cofactor-annotation information becomes available.

## Author Contributions

Sanjay R: conceptualisation, methodology, software, experiments, writing – original draft. B. Rajendra: methodology, supervision, validation, writing – review and editing.

## Data and Code Availability

The dataset used in this study is the publicly available CLEAN benchmark [1], accessible at https://github.com/tttianhao/CLEAN. All feature-extraction code, model-training code, and the exact result files used to generate every number reported in this paper are provided with this submission (`run pilot full.py`, `run pilot v2.py`, and the ESM-2 embedding/classification scripts used to produce Section 3.5) and will be made available at https://github.com/sanjuz-cas/ENZYKAN.

